# Simultaneous Discovery of Positive and Negative Interactions Among Root Microbiome Bacteria Using Microwell Recovery Arrays

**DOI:** 10.1101/2020.01.03.894477

**Authors:** Niloy Barua, Ashlee M. Herken, Kyle R. Stern, Sean Reese, Roger L. Powers, Jennifer L. Morrell-Falvey, Thomas G. Platt, Ryan R. Hansen

## Abstract

Understanding the consequences of microbe-microbe interactions is critical in efforts to predict the function of microbiomes and to manipulate or construct communities to achieve desired outcomes. The investigation of these interactions poses a significant challenge - in part due to the lack of suitable experimental tools. We present the Microwell Recovery Array, a high throughput approach designed to rapidly screen interactions across a microbiome and uncover higher-order combinations of strains that either inhibit or promote the function of a GFP-producing focal species. One experiment generates 10^4^ unique microbial communities that contain a focal species combined with a unique combination of previously uncharacterized cells from plant rhizosphere. Cells are then sequentially extracted from individual co-culture wells that display highest or lowest levels of focal species function using a novel high-resolution photopolymer extraction system. Microbes present are subsequently identified and the putative interactions are validated. Using this approach, we screen the *Populus trichocarpa* rhizosphere for bacterial strains affecting the survival and growth of *Pantoea* sp. YR343, a plant growth promoting strain isolated from the *P. trichocarpa* rhizosphere. We were able to simultaneously isolate and validate multiple *Stenotrophomonas* strains that antagonize strain YR343 growth and a set of *Enterobacter* strains that promote strain YR343 growth. The latter demonstrates the unique ability of the platform to uncover multi-membered consortia that generate emergent phenotypes. This knowledge will inform the development of beneficial consortia that promote the production of *Populus* biofuel feedstock, while the platform is adaptable to screening higher-order interactions in any microbiome of interest.

**Significance Statement:** Achieving a fundamental understanding of microbe-microbe interactions that occur within microbial communities is a grand challenge in microbiology due to the limited experimental tools available. In this report, we describe a new tool that enables one to screen microbial interactions across thousands of compositionally unique communities to discover collections of bacteria that antagonize or promote the survival and growth of bacteria with important functions. This approach has the unique ability to uncover higher-order combinations of bacteria that generate emergent phenotypes, information useful for development of biofertilizer, biocontrol, or probiotic consortia, as well as in the design of communities for biosynthetic compound production.

## Introduction

Microbial communities are often highly diverse and have widespread impacts on human health (1,2), agricultural productivity (3,4), energy production (5,6), and water quality (7). Interactions among the species and strains that co-occur within microbiomes often influence their function and the establishment and success of functionally important taxa (3,8). While genomic and metagenomic approaches have transformed our ability to determine community composition and species co-occurrence patterns (9), understanding how interactions amongst strains impact community structure and function remains difficult. Despite this knowledge gap, there is a considerable need for understanding how natural community structure influences function, how communities respond to environmental pressures, and how communities can be constructed for engineered outcomes. Engineered communities have provided ground-breaking approaches in a few applications such soil clean-up (10) and digestion of municipal solids (11), however the limited understanding of microbial interactions has impeded the use of synthetic communities in the majority of applications.

The high species diversity of many microbiomes necessitates new screening tools that are designed to explore the vast number of potentially important microbe-microbe interactions. These tools must connect an observed cellular or community phenotype with genetic information from the interacting species as well as information on the interaction itself. Classical microbiological techniques for probing interactions rely on manually pairing isolates together (12), an inherently low-throughput approach that in practice is often based on qualitative observations of bulk populations. Micro- and nanoscale devices offer vast improvements by providing high-throughput measurement, observation of single cell behavior, and precise design and manipulation of the microenvironment. These approaches have advanced our understanding of microbial mutualism (13), community adaptation to environmental pressures (14-16), and the role of spatial structure in driving community phenotypes (17-19), among other findings. Recently, Kehe *et al.* (20) introduced the k-Chip, an innovative microscale platform designed to screen multi-membered communities consisting of various combinations of known isolates for emergent phenotypes. While these tools are expected to provide important advancements in our understanding of microbiomes, they are widely limited to *in situ* measurements. Consequently, cells must be identified and manipulated during or prior to the screening observations, which greatly constrains both the number of strains that can be considered and undermines the ability to discover interactions involving unknown strains present in a microbiome.

Here, we present the microwell recovery array (MRA), a discovery-driven tool designed to first screen interactions across thousands of unknown environmental isolate mixtures taken from plant root microbiomes, then uncover pair-wise or multi-species communities that best antagonize or promote the function (e.g. growth) of a non-model focal species (**Fig. *1***). Our strategy relies on a well-characterized stochastic seeding process (21) to randomly combine the focal species — typically one with a known beneficial function (e.g. plant growth promotion) or deleterious function (e.g. pathogenesis) — with a unique combinations of cells from a microbiome into an array of microwells. A 10 μm well diameter was chosen to confine interacting cells together at length scales similar to those found in multi-species biofilms (22), confinement at these length scales often facilitates inter-cellular interactions (23). Cells are then trapped within the wells using a previously developed photodegradable polyethylene glycol (PEG)-based membrane (24), co-cultured, and then focal strain growth in each well is tracked with time lapse fluorescent microscopy (TLFM). Cellular communities showing a desired phenotype (e.g. highly enhanced or diminished focal species growth) can be extracted from any individual well using a patterned light source to spatially ablate the membrane, releasing cells into solution for recovery. The extraction and recovery capabilities are the key enabling features of the platform, allowing for sampling of a microbial community from any number of individual microwells that indicate a desired outcome, in a sequential fashion. Ultimately, this allows one to identify the interacting strains after the screening step, thus thousands of unique species can be included in a single screen. Extraction also enables follow-up phenotypic characterization with standardized assays to confirm the interaction.

**Fig. 1:**
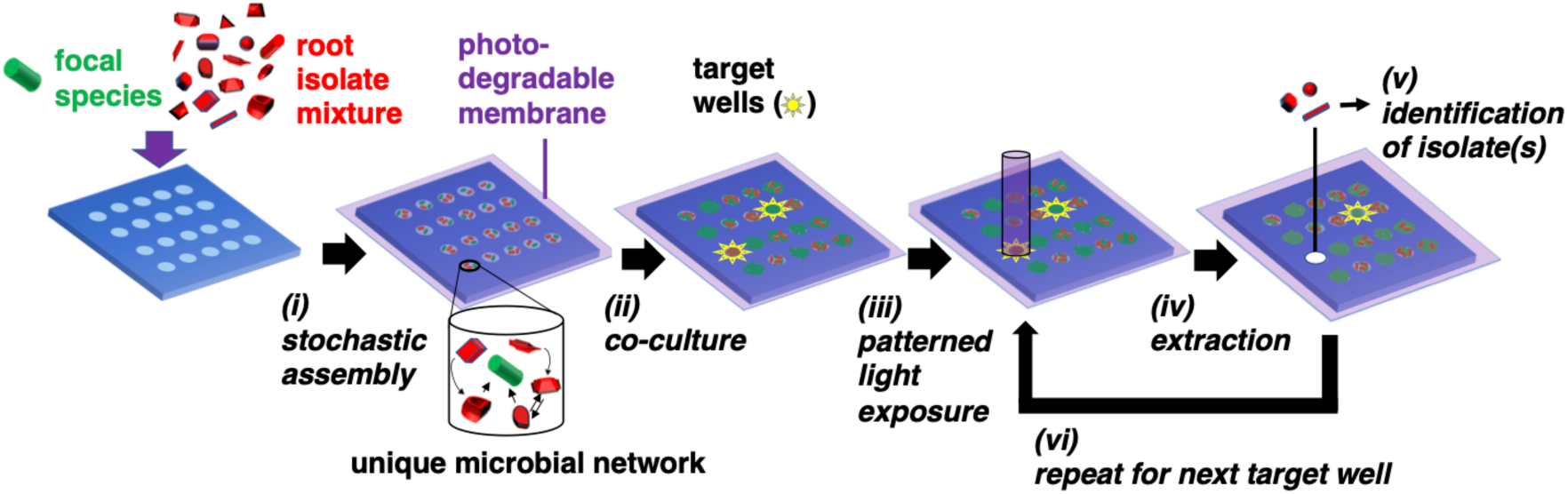
MRAs for screening microbe-microbe interactions. (i) GFP-expressing focal species are combined with a random combination of bacteria cells from an environmental microbiome in a stochastic seeding process. Different shapes represent unique microorganisms. (ii) Cells are trapped within their wells using a photodegradable PEG hydrogel membrane and monitored in parallel during co-culture using TLFM. (iii) The membrane is ablated over a target well showing highest or lowest levels of focal species growth using patterned light exposure, then (iv) isolates are extracted and recovered from an opened well. (v) Isolates are identified using 16S amplicon sequencing. (vi) Steps (iii-v) are repeated in iterative fashion to remove each community of interest.

To develop the approach, we first investigate co-culture in the MRA format using a well-characterized interaction between *Pseudomonas aeruginosa* and *Agrobacterium tumefaciens* (25,26), followed by a microbiome screen using *Pantoea* sp. YR343 as the non-model focal species. Strain YR343 is a Gram-negative, plant growth promoting bacteria (PGPB) isolated from the rhizosphere of an eastern cottonwood *Populus trichocarpa* tree (27,28). As *P. trichocarpa* is a promising biofuel feedstock (29), uncovering interactions that influence the function of beneficial organisms in its rhizosphere has received intensive interest in recent years (30-32). *Pantoea* sp. YR343 can also colonize *Triticum aestivum* and stimulate lateral root formation (27,28). Likewise, related *Pantoea* strains have garnered interest for antibiotic production (33, 34), bioremediation and waste recycling (35, 36), and cancer treatment (37, 38). On the other hand, other *Pantoea* sp. can be pathogenic in plant, animal and human systems (39). Thus, uncovering unique sets of organisms that both promote or inhibit *Pantoea* growth, as demonstrated here, has use in several contexts.

## Results and Discussion

### Microwell Recovery Arrays enable parallel monitoring of microscale co-culture sites and generation of outlier wells with unique growth phenotypes

Our prior results demonstrated that MRAs could be used for parallel tracking of the growth of *P. aeruginosa* PAO1 communities during mono-culture in microwells, where small (5 and 10 µm diameter) wells were used to generate high variations in inoculum densities across the array during the seeding step, and growth outcomes were dependent on inoculum density and the level of spatial confinement present (21). To develop the MRA for multi-species co-culture, here we added *Agrobacterium tumefaciens* C58 to this system. Strains PAO1 and C58 have a well-characterized, competitive interaction *in vitro*, where PAO1 tends to outcompete C58 due to quorum sensing-regulated growth rate and motility advantages (25,26). A 1:100 mixture of C58 cells expressing green fluorescent protein (GFP; hereafter C58-GFP) and PAO1 cells expressing mCherry protein (hereafter PAO1-mCherry) were inoculated into 10 µm diameter wells at a seeding concentration (OD_600_ = 0.1) that generates 20 ± 5 cells per well based on prior characterizations. Under these conditions, PAO1-mCherry and C58-GFP cells are paired together at a high-dispersity due to the stochastic, Poisson seeding process (21).

The MRA platform enabled parallel tracking of species growth according to the respective fluorescence emission signals from each addressable well during co-culture (**Fig. *2A***). End-point growth levels as well as signature growth profiles could be attained from each well. Comparison of end-point fluorescence signals after a 36 hr co-culture period identified that the majority of the wells (96%) generated outcomes where PAO1-mCherry outgrew C58-GFP **(Fig. *2B,C*)**, likely because of a favorable PAO1 seeding ratio and competitive PAO1 growth advantages. However, in the MRA format, outlier testing identified a minority (4%) of wells with communities dominated instead by C58-GFP cells after co-culture (**Fig. *2B,D***). This finding reveals that high-dispersity microbe pairing between competing species produces wells with rare growth outcomes after co-culture. Here, despite C58 cells being present at lower concentrations in the seeding solution, the stochastic seeding process generated a minority of wells with conditions favoring C58-GFP growth. This finding was leveraged towards more complex co-culture systems, where the stochastic seeding and parallel growth tracking features of the MRA are applied to screening interactions in environmental microbiomes.

**Fig. 2:**
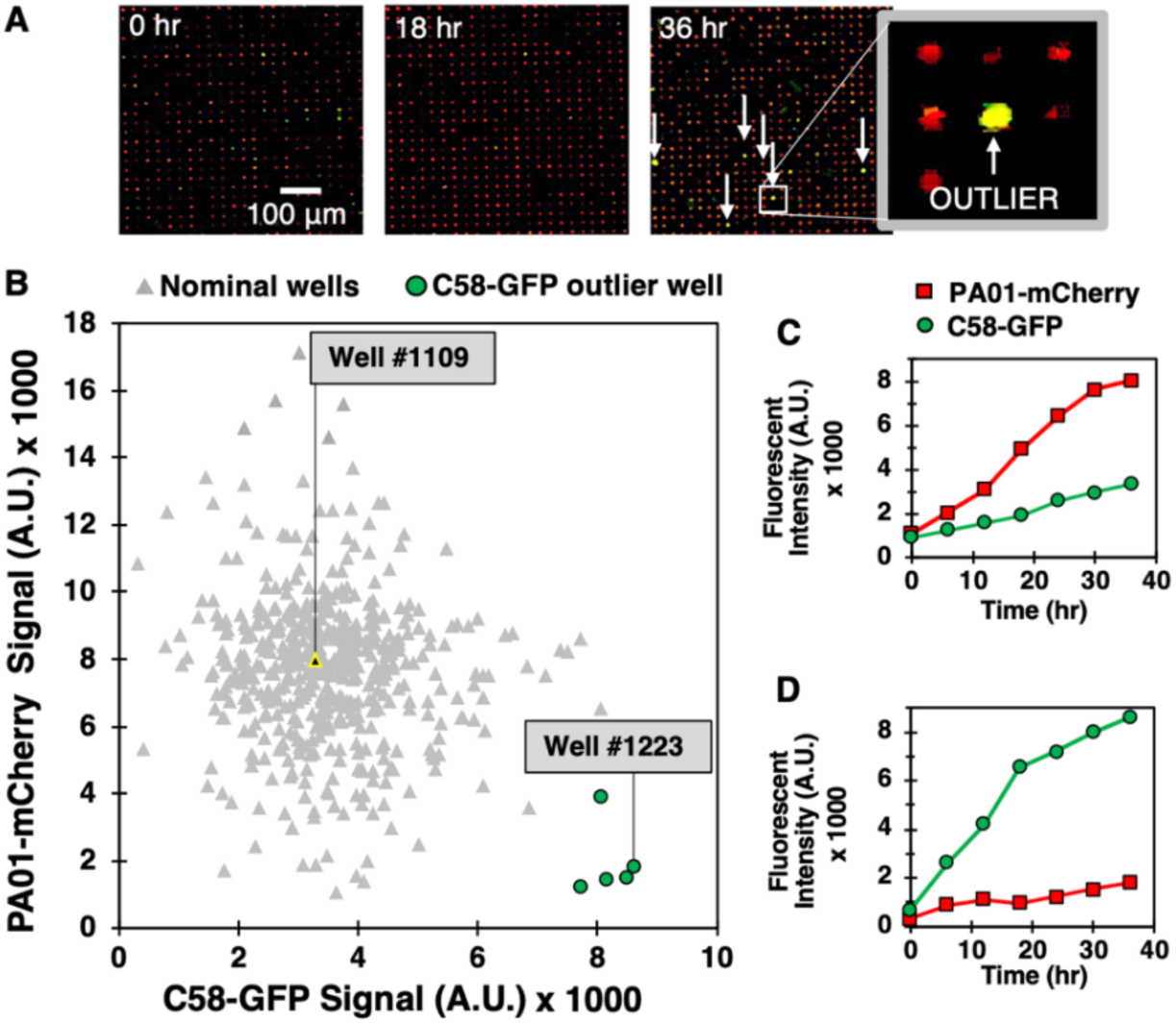
*(A)* Model C58-GFP (green) – PAO1-mCherry (red) co-culture in the MRA. Arrows denote rare outlier wells where C58-GFP outgrew PAO1-mCherry. *(B)* Scatter plot of endpoint (36 hr) green (C58-GFP) versus red (PAO1-mCherry) well signals from a sample 549 well array, with outlier wells identified (green). *(C)* PAO1 growth rate was significantly higher than C58 in nominal wells (p-value<0.0001). *(D)* A significant difference in growth trends were observed in outlier wells, where C58 outgrew PAO1 (p-value<0.0001).

### Co-culture of *Pantoea* sp. YR343 with stochastically assembled communities from the *P. trichocarpa* rhizosphere simultaneously generates positive and negative YR343 growth outcomes

To extend the microwell platform capabilities to screening non-model test species against unknown isolates, we screened rhizobiome samples from the *P. trichocarpa* root microbiome for effects on the growth of focal species *Pantoea* sp. YR343 expressing GFP (hereafter denoted YR343-GFP). Here, we used stochastic seeding, attachment of the photodegradable PEG membrane (24), and focal species growth monitoring to identify rare combinations of cells generating unique YR343-GFP growth profiles (**Fig. *1***, steps i and ii). Cell mixtures were first seeded into 10 µm diameter microwells at high density (OD_600_ = 0.2) and at a 1:100 YR343:isolate ratio, cultured, and growth kinetics in each well were tracked over the course of 12 hrs using TLFM. For comparison, monoculture arrays consisting of only YR343-GFP focal species were also evaluated. In each case, YR343-GFP growth was evaluated in 225 co-culture microwells from a 15×15 well grid (**Fig. *S1***) across 16 selected arrays on a single substrate (n=3600 microwells total). Here, R2A media was chosen as a generalist culture media. This media has been used to recover more than 300 phylogenetically diverse isolates from *P. trichocarpa* rhizosphere and endosphere samples, and so should permit co-culture of a large number of combinatorial strain mixtures within the microwell environment (40, 41). While the YR343-GFP monoculture generated growth profiles across the array with relatively low variance (σ^2^=3.55) according to final end-point fluorescence levels, mixed cultures generated a wider range of growth profiles, with final growth levels of higher variance (σ^2^=17.55), indicating an impact due to the addition of the environmental isolates (**Fig. *3A-C***).

**Fig. 3:**
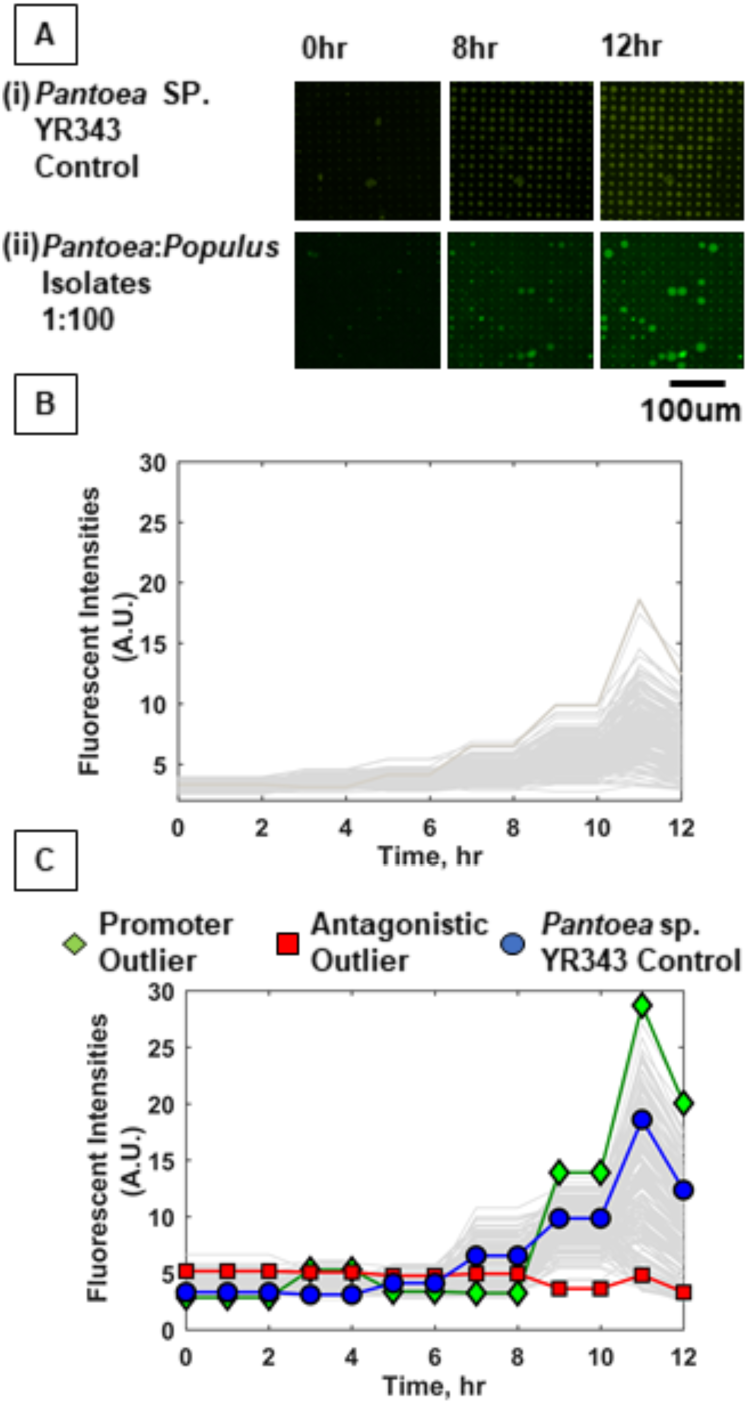
YR343-GFP growth in mono-culture and co-culture within 10 µm microwells. *(A)* TLFM images of a sample 15×15 array of microwells after (i) seeding only YR343-GFP or (ii) seeding YR343-GFP with isolates from a *P. trichocarpa* rhizobiome. *(B)* Growth curves generated from a sample 900 microwell array during YR343-GFP mono-culture, or *(C)* YR343-GFP co-culture with rhizosphere isolates. Outlier wells representing growth promoting and antagonistic communities, respectively were identified from the growth curves.

In co-culture, 14% of the wells contained microcolonies that appeared to grow out of the wells and into the membrane space, causing a microcolony diameter to expand beyond the well diameter (>10 µm) (**Fig. *3Aii***), indicating a positive interaction. While the locations on the array where this effect occurred appeared random, we checked for the possibility of crosstalk between neighboring wells, where a developing microcolony may influence growth in another well due to diffusion of metabolites or other biomolecular products. Of wells with this enhanced growth phenotype, 2% of neighboring wells also showed this phenotype, suggesting that well-to-well crosstalk can occur. On the other end, wells showing decreases in well fluorescence signal were also identified, these decreases were caused by lysis of the focal species and GFP diffusion from the wells, as previously characterized when using PAO1 as the focal species (21). This effect was noted in 34% of co-culture microwells. Wells showing the highest decrease in fluorescence signal were identified as containing candidate cells antagonistic to YR343-GFP.

### Sequential extraction, recovery and identification of isolates from microwell communities

Following on-chip analysis in mixed culture arrays, the patterned illumination tool was used to extract communities from the five wells with highest fluorescence signal after 12 hours of culture (**Fig. *4A,B***). This was followed by extraction of communities from four wells with the lowest levels of YR343-GFP growth, and four wells where YR343-GFP grew to intermediate levels. Extraction required exposure of a patterned 365 nm light (20 mW/cm^2^, 10 min) to remove the membrane over the well following the approach in van der Vlies *et al.* (24). Membrane removal was confirmed by bright-field microscopy, at which point cellular material was observed moving out of the wells and into solution (**Fig. *4B***). After exposure, arrays were washed with extraction buffer (R2A media + 0.05% Tween20 solution) to retrieve cells from an opened well. Extraction buffer was then plated onto R2A-agar for growth and recovery of individual colonies. During our previous characterizations of this procedure, we noted that >99.9% of bacteria originated from opened wells as opposed to outside contamination (24), which provided high confidence that the recovered product here originated from the target well. We also previously observed that bacteria could be completely removed from wells after washing (24), thus we expected minimal cross-contamination when opening additional wells for further sampling. After recovery, phylogenetic analysis based on 16S rRNA sequences of all strains isolated from each targeted microwell was performed (**Fig. *S2*)**. The analysis revealed that all extracted microwells identified as growth promoting for YR343-GFP (5 of 5) harbored *Enterobacter* sp. / *Pantoea* sp. strains, and one of these wells contained at least one *Pseudomonas* sp. strain. In stark contrast, all wells identified as antagonistic to YR343-GFP contained at least one *Stenotrophomonas* sp. strain. Several of these wells (3 of 4) also contained at least one *Enterobacter* sp. or *Pantoea* sp. strain (**Fig. *4C***).

**Fig. 4:**
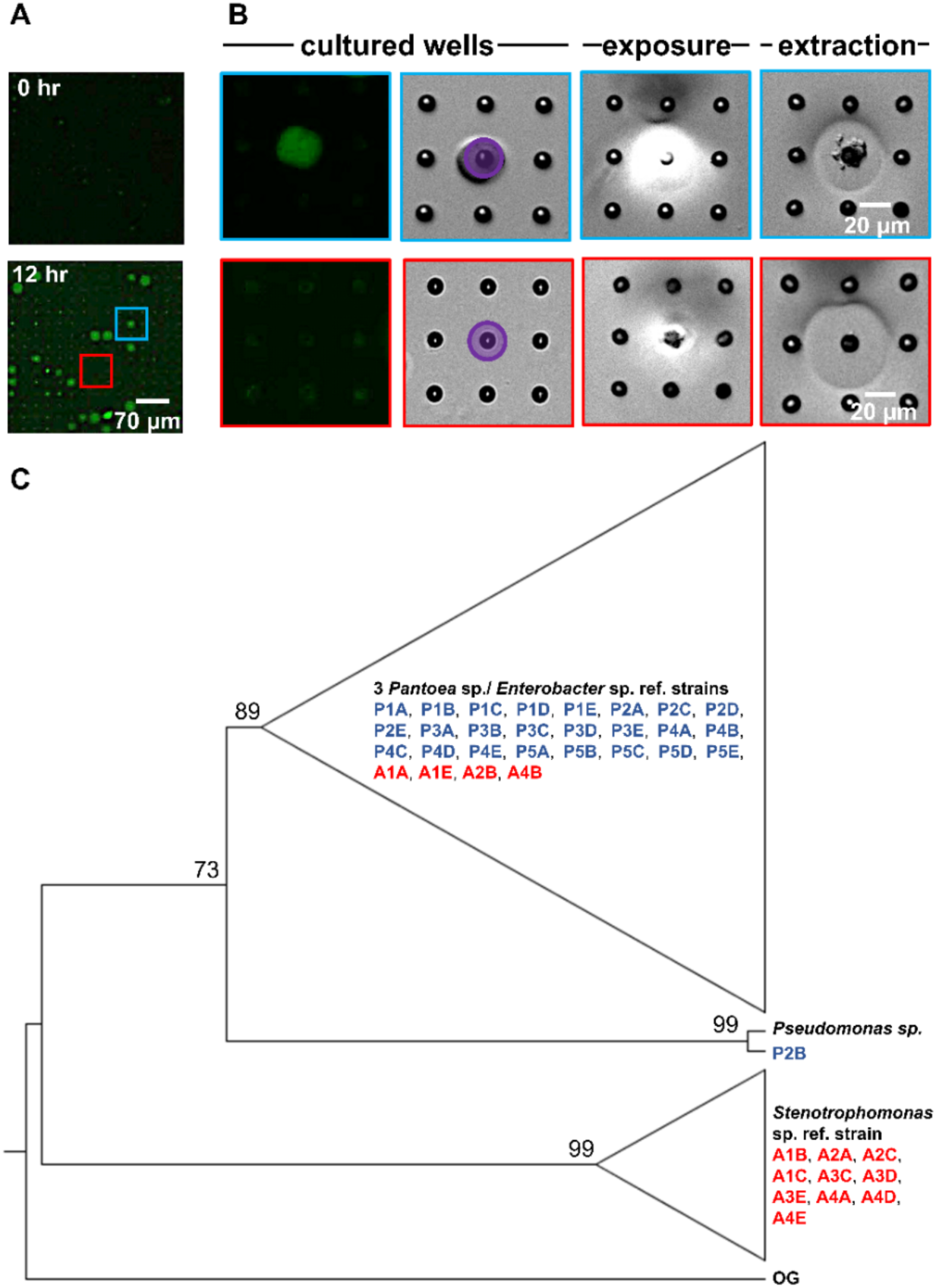
Sequential removal of growth-promoting and antagonistic communities from an array sub-section after co-culture. (*A*) Microwell array before and after co-culture. This 15×15 microwell array contained both a YR343 growth promoting community (blue) and YR343 antagonistic (red) community that were targeted for extraction. (B) Targeted removal of the microwell community in which YR343 grows to its highest observed end-point fluorescence (top row, blue outline), followed by targeted removal of a microwell community in which YR343-GFP grew poorly (bottom row, red outline). Purple area denotes UV exposure area used for membrane degradation. (C) Maximum likelihood phylogenetic tree based on partial 16S rRNA sequences (921 sites) of select reference strains and isolates extracted from promoted (P) and antagonized (A) wells. We collapsed the branches of the monophyletic group composed of *Enterobacter* sp. and *Pantoea* sp. strains and the clade of *Stenotrophomonas* sp. strains. *A. tumefaciens* C58 was used as the outgroup (OG) organism and the following reference strains were included: *Pantoea* sp. YR343, *Enterobacter cloacae* E3442, *Enterobacter ludwigii* KUDC1774, *Pseudomonas putida* S13.1.2, *Stenotrophomonas maltophilia* NCTC10259. We labelled nodes with corresponding bootstrap percentages.

### Interactions can be recapitulated in 96-well plate format for validation

The extraction of cellular communities from the MRA allows for off-chip validation and characterization of the microbial interactions observed during the screen. This capability is critical for validation, as the high density of microwells (625 wells/mm^2^) has potential to cause well-to-well cross-talk due to diffusion of molecules across the array, a factor that may generate false-positive outcomes. To address this, we investigated how the strains recovered from targeted microwells influenced the growth of YR343-GFP in 96-well plate format. This represented a scaled-up environment (from 1.6 pL microwell volumes to 100 µL solution volumes) that precludes diffusive crosstalk from neighboring wells.

For these evaluations, we hypothesized that both growth promotion and inhibition measured in MRA format resulted from diffusive interactions between the focal species and the collection of isolates present within a well. To test this hypothesis, YR343-GFP was cultured in 96-well plate format in media conditioned by four selected isolates recovered from a selected antagonist well (Well A4, **Fig. *S2***). Conditioned media was obtained by first culturing isolates in R2A media to stationary phase, then removing the cells to obtain cell free culture fluid (CFCF). Fresh R2A media was then added to the CFCF in a 1:1 volumetric ratio to supply growth nutrients, and YR343-GFP was inoculated for growth monitoring. Conditioned media obtained using CFCF from a combined co-culture of all 4 antagonistic strains was also evaluated. These growth curves were compared to YR343-GFP growth in unconditioned media, which consisted of R2A media instead supplemented with blank 1X PBS buffer at the same volumetric ratio. Growthcurver R was then used to estimate bacterial carrying capacity and growth rate (42) in each experiment (**Fig. *S3***). Congruent with microwell observations, we observed that conditioned media from 4 isolates significantly reduced the carrying capacity and growth rate of YR343-GFP compared to its culture in unconditioned media (**Fig. *5A***). Conditioned media from the combined 4-member antagonist combination also showed significantly lower carrying capacity and growth rate compared to the unconditioned control media. CFCF from *Stenotrophomonas* isolate A4A had statistically equivalent growth metrics as that from the CFCF consortia, suggesting that this strain is the most potent inhibitor of YR343.

To investigate the effect of the strains identified as YR343 growth promoters, YR343-GFP growth was again monitored in media conditioned with CFCF from clonal cultures, here using 5 isolates selected from a selected promoter well (Well P3, **Fig. *S2***). To test for an emergent effect, conditioned media containing CFCF produced from a co-culture of the combined 5 isolates was also used. When YR343 growth in conditioned media from the CFCF of individual isolates was measured, only two were able to increase growth rate and one was able to increase carrying capacity (**Fig. *5B*, Fig. *S4***). Strikingly, however, the CFCF from the 5-member consortia was able to provide highest increases in both YR343 growth rate and carrying capacity. The 5-member consortia also provided a statistically significant increase in carrying capacity compared to isolate P3B, the individual isolate that generated the highest increase in YR343 carrying capacity after conditioning media on its own. These findings indicate that the observed enhancement of YR343’s population growth depends on the presence of multiple strains, and is not simply the consequence of a pairwise interaction. As such, the enhanced YR343 growth is an emergent property of the community of species recovered using the MRA, demonstrating the power of this approach to identify functions dependent on higher-order interactions among bacterial species.

**Fig. 5.**
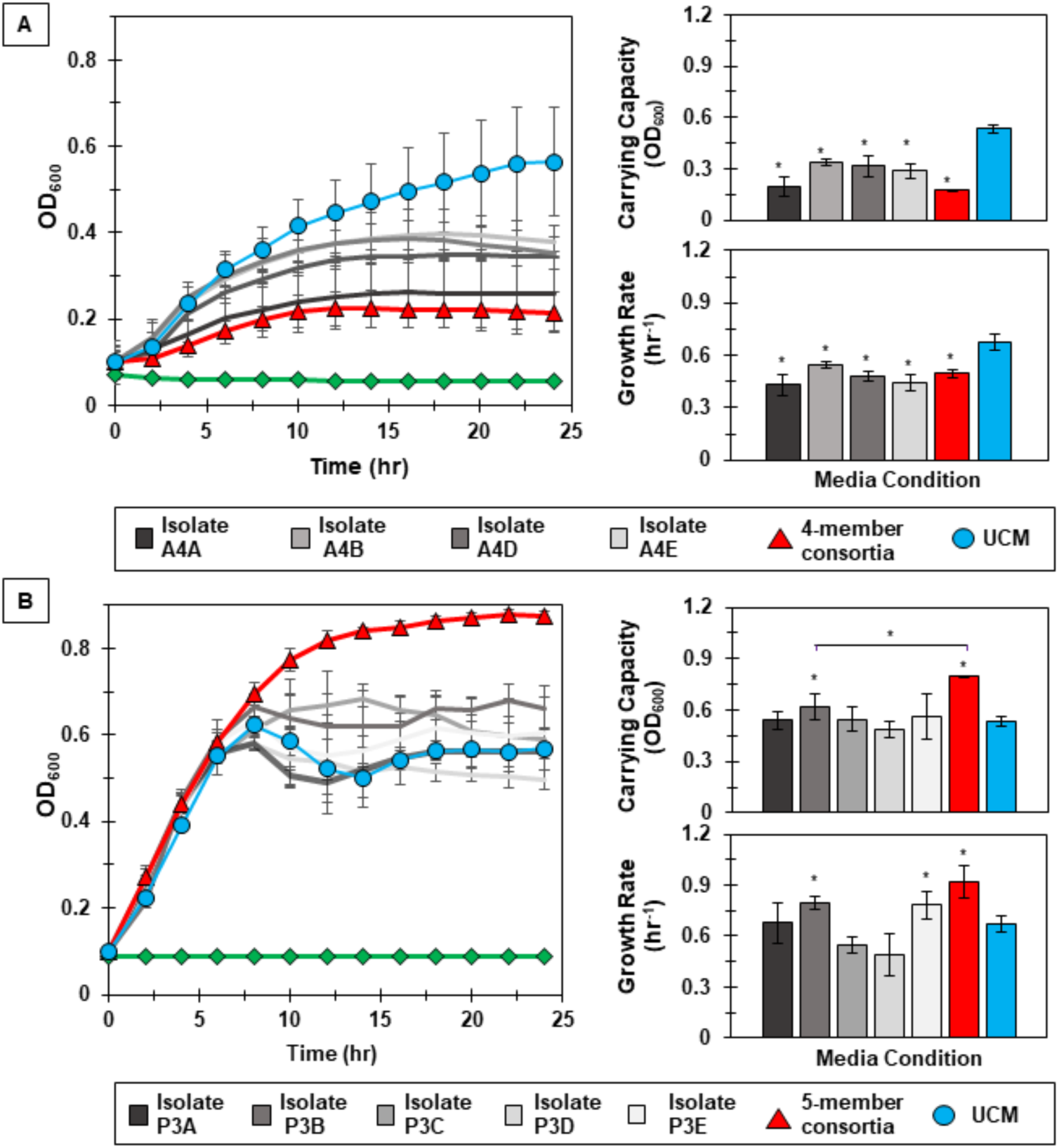
Interactions identified in the MRA can be validated in 96-well plate format. *(A)* Left: YR343 growth curves after inoculation into conditioned media from the antagonistic isolate, the isolate consortia, or unconditioned media (UCM). The control (green line) is conditioned media that was not inoculated with YR343 to verify that there was no growth carry over or contaminating microbes present. Right: Corresponding carrying capacity and growth rates for each growth curve. *(B)* Left: Analogous YR343 growth curves after inoculation into conditioned media from a promoter isolate or the promoter isolate combination. Right: Corresponding carrying capacity and growth rates. All growth experiments occurred at 28°C, 215 RPM. Statistical differences were identified by comparison of growth metrics between YR343 culture in conditioned media from each isolate or isolate mixture and YR343 growth in UCM (Wilcoxon two-sample test, *=P<0.01, n=6 independent experiments).

## Conclusion

The MRA examines thousands of unique, multi-species microbial communities to discover both antagonistic and growth promoting interactions on a focal species. Using this new approach, we simultaneously identified strains that antagonize focal species growth, as well as multi-strain consortia that promote focal species growth only when co-cultured in combination. The platform is the first of its kind, unique because it (i) screens organisms that are unknown during the screening step, dramatically expanding the number of interactions and cellular combinations that can be accommodated, and (ii) screens in combinatorial fashion to uncover higher-order microbial networks that generate emergent phenotypes. The key innovation underlying this capability is the ability to recover cells from specific microwells of interest, thereby allowing for subsequent off-chip genetic characterization for species identification then phenotypic characterization for validation of the interaction. The current format also does not require a microfluidic setup, often a technical barrier for implementation into standard microbiology labs (43).

For the first generation of the MRA, we have developed its use towards screening interactions that influence growth phenotypes, however the approach is amenable to screening microbiomes for microbes that have positive or negative effects on other focal species functions, provided that the function can be coupled to a fluorescence reporter (e.g. a GFP promoter-reporter). This may include microbial interactions that affect quorum sensing activation (44), virulence factor expression (45), and plasmid conjugation (46), to name a few. While demonstrated here for the *P. trichocarpa* root microbiome, the platform is directly amenable to screening interactions across any microbiome where high species diversity is present, which may include the gut, the soil, freshwater and marine ecosystems, and other rhizosphere environments.

## Materials and Methods

### Bacteria strains and P. trichocarpa samples

Bacteria strains and plasmids used are listed in **Table *S1***. We introduced pSRKKm-sfGFP into *A. tumefaciens* C58 and pSRKKm-mcherry into *P. aeruginosa* PAO1 via mating with *Escherichia coli* S17-1 λpir carrying the respective plasmids using previously described methods (47). These plasmids were transformed into competent S17-1 λpir *E. coli* strains using calcium chloride heat-shock transformation. *Pantoea* sp. YR343-GFP constitutively expresses EGFP from a chromosomal insertion as previously described by Bible *et al.* (27). All strains and isolates used were stored in 25% glycerol at −80 °C.

### MRA design and fabrication

MRAs were designed to contain 10 μm diameter microwells etched to 20 μm well depths, spaced at a 30 µm pitch. The array consisted of a 7×7 grid of sub-arrays, each sub-array contained a 15×15 array of microwells, totaling 11,025 microwells available for analysis. Each well in the 15×15 sub-array was assigned with its own unique on-chip address for identification using brightfield microscopy (**Fig. *S1***). Microwell arrays were fabricated on 3-inch diameter N-type silicon wafers (University Wafers) after coating with a 1μm thick layer of Parylene N (PDS 2010 Labcoater, Specialty Coating Systems). Arrays were then fabricated in a cleanroom environment using photolithography (**Fig. *S5***) following previous protocols (21).

### Bacteria seeding and trapping on microwell arrays

C58*-*GFP and PAO1-mCherry were grown in LB and YR343-GFP was grown in R2A media to mid-log phase and then resuspended in their respective growth media to an OD_600_ of 0.2. To inoculate microwell substrates, 700μL of this cell suspension was then incubated over the substrates at room temperature for 1hr **(Fig. *S6*)**. The substrates were dried and the parylene was peeled off of the microwell surface along with the cells attached to the background regions of the array by applying Scotch tape and forceps (21). For studies involving C58-GFP and PAO1-mCherry co-culture, the seeding solution contained C58-GFP and PAO1-mCherry cells in a ratio 1:100 and a total OD_600_ of 0.1. For studies involving YR343-GFP and *P. trichocarpa* rhizobiome co-culture, washed YR343-GFP cells and *P. trichocarpa* rhizobiome cells were mixed to achieve a YR343-GFP:isolate ratio of approximately 1:100 in the seeding solution at an OD_600_ of 0.2. To keep the cell concentrations of C58-GFP in both mono-culture and co-culture experiments constant, PAO1-mcherry at OD_600_=10 was added to C58-GFP at OD_600_=0.1 to reach a C58-PAO1 ratio of 1:100. The inoculum was then diluted to OD_600_=0.1 and 700µm of this inoculum was then seeded into microwell array substrates as described above. Similarly, OD_600_=0.2 cultures of *P. trichocarpa* rhizobiome was mixed with OD_600_=20 of YR343-GFP to reach a YR343-*P. trichocarpa* ratio of 1:100. This seeding suspension was diluted to OD_600_=0.2 and seeded on top of microwell arrays for co-culture studies.

### Time lapse fluorescence microscopy (TLFM)

A Nikon Eclipse Ti-U inverted microscope with NIS Elements software, a motorized XYZ stage, a humidified live-cell incubation chamber (Tokai Hit), and a DS-QiMc monochromatic digital camera was used for TLFM measurements. Seeded microwell arrays sealed with either a LB-agar coverslip or the photodegradable membrane (**Fig. *S6***) were placed in a custom 3D printed scaffold designed to accommodate the microwell array while submerged under liquid media. The scaffold aided in image acquisition by maintaining a constant distance (100 µm) between the array and the glass slide, enabling the microwell substrate to stay within the focal plane during the culture period (**Fig. *S7***). The scaffold along with the inverted microwell substrate were placed inside a humidified live-cell incubation chamber at 28°C during imaging. A FITC filter was used to image C58-GFP strains (20×, 200 ms, 17.1× gain) and a TRITC filter was used to image PAO1-mCherry strains (20×, 300 ms, 17.1× gain). For YR343-GFP, images were taken with a FITC filter (20×, 300 ms, 36× gain) with a neutral density filter with 25% standard light intensity to ensure imaging without photobleaching. Brightfield images were also taken at each section of the array after fluorescent imaging. Images of the microwell arrays were taken every 60 minutes during culture. Green and red fluorescent images from the C58-GFP and PAO1-mCherry co-culture system were analyzed using Protein Array Analyzer tool in ImageJ to generate growth profiles for each organism. YR343-GFP in monoculture or mixed culture was evaluated using an image analysis routine in MATLAB to identify wells with highest and lowest growth levels for extraction.

### Recovery of isolates from wells and identification with 16S RNA sequencing

The extraction procedure was slightly modified from van der Vlies *et al.* (24) and is described in Supplementary Information. Following extraction, extract containing the suspension of cells from an individual microwell was plated onto R2A media. Five colonies were sampled from each extracted microwell based on differences in colony morphology and color. Individual colonies were cultured in R2A media and genomic DNA of each isolate was extracted using the Promega (Madison, WI) Wizard ® DNA Purification kit and sequenced at Genewiz (South Plainfield, NJ, USA) for 16S ribosomal RNA (rRNA) sequencing of the V1 to V9 regions. The sequences were aligned using MUSCLE (48) and generated a maximum likelihood phylogenetic tree based on partial 16S rRNA sequences (921 bp) using PhyML 3.0 (49) with 1000 bootstrap replicates and using the Smart Model Selection (50) tool based on Akaike Information Criterion, a starting tree estimated using BIONJ, and the NNI method for tree topology improvement (**Fig. *4C*, Fig. *S2***).

### Validation using 96 well plates

To obtain CFCF from individual isolates, each isolate was cultured (28°C, 3000 rpm) in 2mL of R2A broth media overnight, and then cells were removed from the media by centrifugation (2000g, 10 min). To obtain CFCF from combinatorial mixtures, isolate panels were instead inoculated together in R2A media and cultured overnight, followed by cell removal by centrifugation. To obtain conditioned media, isolate CFCF was mixed with YR343-GFP in fresh R2A media at a 1:1 volumetric ratio to reach an initial OD_600_ value of 0.1 (final volume = 100 µL), at which point growth was quantified with a Biotek Epoch 2 Multi-Mode Microplate Reader (28°C, 300rpm). Unconditioned media was obtained following the same procedure except 1X PBS was added to fresh R2A media instead of isolate CFCF. To verify the OD_600_ measurement was due to YR343-GFP growth, CFCF from selected isolates without inoculation of YR343 was also measured. A total of n=6 independent replicates were measured for each culture condition. Growth rates and carrying capacities of each condition were quantified using Growthcurver and compared using the Wilcoxon two-sample test.

## Notes

To date there are no competing financial conflicts of interest to declare.

## Acknowledgments

We acknowledge support from the National Science Foundation (MCB-1650187) and the U.S. Department of Energy (DE-SC0018579). A portion of this research was sponsored by the Genomic Science Program, U.S. Department of Energy, Office of Science, Biological and Environmental Research, as part of the Plant Microbe Interfaces Scientific Focus Area (http://pmi.ornl.gov). Oak Ridge National Laboratory is managed by UT-Battelle LLC, for the U.S. Department of Energy under contract DE-AC05-00OR22725. We thank Priscila Guzman for making the C58 and PAO1 strains used in experiments. Nanofabrication was conducted at Nebraska Nanoscale Facility at University of Nebraska Lincoln.

