## Supplementary material for "Simultaneous Discovery of Positive and Negative Interactions Among Root Microbiome Bacteria Using Microwell Recovery Arrays": Supplimentary Information

***P. aeruginosa* and *A. tumefaciens* culture.** A model system comprised of *Agrobacterium tumefaciens* C58 expressing GFP (C58-GFP) and *Pseudomonas aeruginosa* PAO1 expressing mCherry (PAO1-mCherry) was used to characterize seeding behavior and interaction in microwell recovery arrays (MRAs). Glycerol stocks were prepared for both strains and stored at -80°C. C58-GFP and PAO1-mCherry were cultured on LB-agar for 24 hrs at 28 and 37°C respectively. For liquid culture of C58-GFP and PAO1-mCherry, a single colony was picked using a sterile inoculation loop, inoculated in 2 mL LB media (10g BactoTryptone, 5g Yeast, 10g NaCl and 15g agar per 1000mL) in sterile test tubes and cultured in a shaker (28 and 37°C respectively, 215 rpm) for 24 hrs. 150 µg Kanamycin and 0.5 µg IPTG (Sigma) were added to each ml of the liquid culture for both C58-GFP and PAO1-mCherry cultures. The tubes were then centrifuged (Eppendorf Centrifuge 5702 Series Ea) and the media was changed every 24 hrs for up to 1 week to keep cells viable.

**Extraction of isolates from *Populus* roots.** A sample of Nisqually-1 *Populus trichocarpa* root was obtained from the greenhouse facilities at Oak Ridge National Laboratory (ORNL). For extraction of isolates, roots were first removed from soil and the aerial parts of the plant were separated from the root system. Large soil aggregates were removed by manually shaking by hand. The remaining portions of the roots were removed with sterile blades. Root pieces were then washed extensively with 1.5 L of sterile ice-cold PBS-tween20 solution (7 mM $Na_2HPO_4$, 3 mM $NaH_2PO_4$, pH 7.0 and 0.05% tween20). The washed solution was filtered through 0.45µm sterile syringe filters (Whatman) to remove larger particles in the suspension. The filtered solution was centrifuged for 15 min at 4400 rpm to obtain the pellet containing rhizosphere-enriched isolates (1). Numbers of glycerol stocks were prepared for *P. trichocarpa* root isolates and stored frozen at -80°C.

**Growth studies of *Pantoea* sp. YR343 monoculture.** For solid phase cultures, *Pantoea* sp. YR343 expressing GFP (YR343-GFP) was cultured in R2A-agar media (pH: 7.2 ± 0.2, Thermo Fisher) at 28°C for 24 hrs. For liquid culture studies, single colonies were picked using sterile inoculation loops and mixed in 2 mL R2A broth media (pH: 7.2 ± 0.2, Teknova) supplemented with Kanamycin (150 µg/mL) in sterile test tubes and cultured for 24 hrs (28°C, 215 rpm). To measure growth parameters of YR343-GFP, liquid cultures were diluted to $OD_{600}$ = 0.1 and 100µL was added to a 96 well plate (28°C, 600 rpm) and absorbance ($OD_{600}$) readings were taken every 10 min. For YR343 monoculture, a lag phase of 2.5 hrs and growth rate of 0.673 $hr^{-1}$were measured, the culture reached stationary phase at 8 hrs (2, **Fig. *S8***).

**Fabrication of LB-Agar coated PDMS coverslips.** To prepare LB-agar coated PDMS coverslips, 21 g PDMS monomer and 3 g of curing agent were mixed for 3 min, degassed for 30 min, then placed in a 6 in. diameter polystyrene petri dish, degassed again for 30 min and baked at 80°C for 2 hrs. PDMS coverslips were ~ 2000 µm thick. Sterile 25×75mm PDMS coverslips were then cut from the dish, placed in a second polystryene dish and coated with LB agar by evenly pouring 3 mL of boiling LB agar over the coverslips and the dish. The dish was cooled at 4°C for 30 min to allow the media to solidify over the coverslips and to ensure minimum dehydration. The thickness of the agar layer was ~100 µm. After seeding the microwell substrate with bacteria, microwells were immediately sealed with the coverslips and placed in the humidified, live cell incubator chamber to conduct growth experiments (**Fig.** ***S6***).

**Photodegradable membrane attachment.** The procedure for attaching the photodegradable hydrogel membranes to microwell arrays is described in van der Vlies *et al.* (3) was used here. 25×25 mm clean glass slides (Fisher Scientific) were first functionalized with a non-reactive silane layer by incubating in 20 mL of 0.5% v/v trichloro(1H,1H,2H,2H-perfluorooctyl) silane in toluene for 180 min (3). Phosphate buffer saline (PBS) with LB solution was then prepared by adding $NaH_2PO_4$ to LB liquid media to reach 100mM final phosphate concentration and adjusting to pH 8 by adding 5M NaOH (aq.). The membrane precursor solution was then obtained by mixing 12.5 µL of PBS-LB solution with 5.6 µL of photodegradable PEG diacrylate monomer (MW 3400) and 6.9 µL of a four-arm PEG thiol solution monomer (MW 10000, NOF America Corporation, DE-100SH, 4). The concentrations of both PEG diacrylate monomer and four-arm PEG thiol solution monomer were 22mM in the precursor solution. 15µL of the liquid pre-cursor solution was then quickly pipetted on top of the perfluoroalkylated glass slides, and the solution was placed over a seeded microwell substrate. Metal spacers were used to provide a constant 38 µm gap between the glass slide and the microwell for the precursor solution. The substrate was then incubated for 25 min at room temperature to allow for membrane formation through monomer crosslinking and gelation (3). The glass slide was then carefully separated from the membrane-functionalized microwell array and the microwell array was placed on top of a PDMS coverslip and added to the 3D printed scaffold for culture and imaging.

**Fabrication of 3D printed scaffolds.** A custom 3D printed scaffold was designed for imaging microwells with time lapse fluorescence microscopy (TLFM). The 3D printed nylon scaffold was designed using Blender software (5). A seeded and membrane-functionalized microwell was placed in the scaffold, the scaffold was then placed over a standard glass microscope slide. The scaffold holds the sealed microwell substrate approximately 100 µm above the slide surface, as shown in **Fig.** ***S7***. This allows for bacteria within the wells to receive nutrients and allows the microwell substrate to remain fixed within the focal plane of the 20× objective, eliminating drift in the x,y, and z-directions during the culture period.

**Image Analysis.** Time-lapse fluorescent microscopy and fluorescence-based image analysis can be used to generate and analyze bacteria growth trajectories in this microwell format, as recently described by Timm *et al.* (6). ImageJ was used to quantify growth trends of the C58-GFP and PAO1-mCherry. MATLAB was to identify wells with highest and lowest levels of growth for YR343-GFP monoculture and co-culture studies. Here, simultaneous brightfield and fluorescence images of each array subsection consisting of 15×15 microwells were taken every hour for a 15 hr culture period. Brightfield and fluorescence images were imported into MATLAB and sorted based on subarray location. The location of the wells was recorded and fluorescence intensities were averaged across each individual well and subtracted from background levels for each time point. Average growth rates and end point well intensities were then quantified across the entire microwell population. Outlier wells with highest levels of deviation in end-point fluorescence (t=12 hrs) were identified as target wells using the Grubb's outlier test (7) and their address was recorded. From these outlier wells, the top 5 growth promoting wells with highest average growth rates and top 4 antagonist wells with the lowest average growth rates were picked for extraction.

**Membrane degradation and well extraction.** Extraction of cell aggregates from microwells followed the protocol recently described in van der Vlies *et al.* (3). An Olympus BX51 upright microscope equipped with an Infinity 3-1 microscopy camera (Lumenera) and Infinity Analyze software was used to identify the location of target wells according to the on-chip well address. Greyscale images of targeted microwells were taken during extraction with a 20×/0.5NA objective. The Polygon400 photo-patterning instrument (Mightex) containing a 365 nm high-power LED source (50 W) was used to project UV light patterns onto the target well locations. The instrument was attached to the BX51 microscope through an adapter containing dichroic filter cube. A BioLED light source control module equipped with a BioLED analog and digital I/O control module was used to control the light source and a liquid light guide was used to deliver light to the Polygon400. Prior to extraction, the Polygon400 was calibrated using PolyScan2 software and a calibration mirror. For extraction, a cultured microwell array substrate with the attached membrane was first submerged in 1mL R2A broth media to prevent membrane dehydration, then placed under the microscope. PolyScan 2 software was then used to define the irradiation pattern, light intensity and irradiation time. Here, a 165×295μm rectangular working area was defined to accommodate an array of microwells. After a targeted microwell was located, it was exposed a 20 μm diameter circular pattern (20 $mW/mm^2$, 10 min) to erode the polymer matrix over the well. The opened microwell array was then washed with R2A broth media + 0.05% Tween20 (5×2mL) to extract cells from the opened microwell. The 10 mL wash solution was centrifuged at 2000g for 10 min and the supernatant was carefully removed leaving approximately 2 mL of solution inside the culture tube.

**Sequential isolate extraction and isolate naming convention.** Extraction occurred from the microwell array in a sequential fashion, first from five different target wells in which YR343-GFP exhibited promoted growth (P1, P2, P3, P4, P5), then four different target wells in which YR343-GFP exhibited antagonized growth (A1, A2, A3, A4). Extracts from each well were plated onto solid R2A media (28°C, overnight) for recovery. After culture, five distinct isolates (A, B, C, D, and E) were picked based on unique colony morphology and streak purified. Colonies were again cultured in liquid media overnight (28ºC, 215 rpm) and stored in glycerol stocks at -80°C. Isolates in **Figs. *4C* and *5*** and **Fig. *S2*** are labeled according to the microwell they were isolated from, then the order at which it was extracted from the array, then the order at which it was picked from the plate after recovery. For example, isolate A4A, the isolate that most strongly antagonizes YR343 growth, was extracted from the fourth antagonistic well and was the first colony picked from the R2A plate.

**96-well plate validation**. Separate CFCF from all 5 isolates were mixed together at equal volumes then added to *Pantoea* cell culture in R2A media at a volumetric ratio of 1:1 to reach an $OD_{600}$ value of 0.1. 100μL of each treatment with 6 independent replicates were cultured overnight in 96 well plates to determine the influence of outlier isolates on the $OD_{600}$ of *Pantoea* sp. YR343 growth. Wilcoxon Two-sample tests were conducted to test whether there is a significant difference between median values of isolate-*Pantoea* combinations and *Pantoea* monoculture (8).

**16S RNA sequencing of isolates**. A single colony of each isolate was cultured, with shaking, overnight in R2A broth. The Gram positive and Gram negative bacteria protocol of the Wizard® Genomic DNA Purification Kit (Promega) was used for genomic DNA preparation. Samples were diluted to 20 ng/μL in 20μL aliquots and the 16S rRNA was sequenced by Genewiz (South Plainfield, NJ). Sequences were assembled using the de novo Geneious assembler (9). A maximum likelihood phylogenetic tree of the isolate 16S sequences and several reference strains was estimated using PhyML 3.0 (10) and visualized using FigTree v1.4.4 (http://tree.bio.ed.ac.uk/software/figtree/). Isolates from A1 – A5 wells are phylogenetically related to either *Enterobacter* sp./ *Pantoea* sp. strains or *Stenotrophomonas* sp. strains. In contrast, most isolates from the P1 – P5 wells belong to the *Enterobacter* sp. + *Pantoea* sp. clade, with only one strain being phylogenetically related to another group of bacteria (**Fig. *S2***).

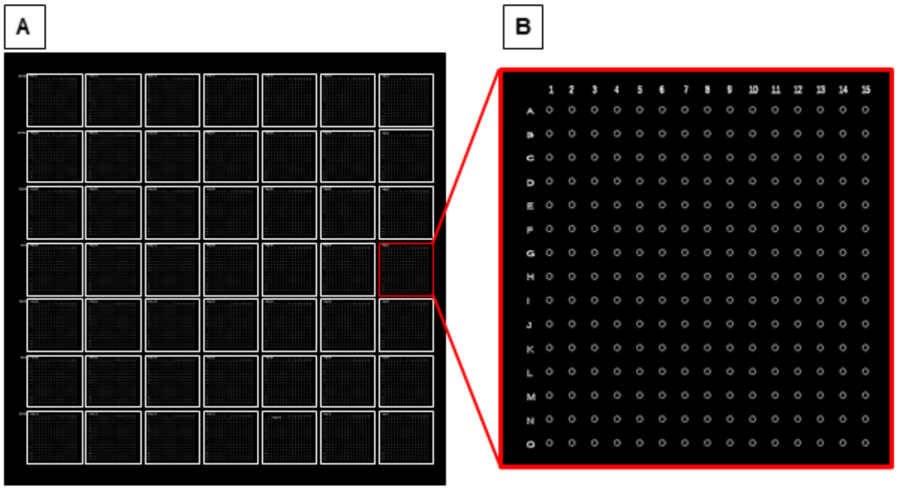

**Fig. *S1*.** MRA layout. (A) The 10μm diameter wells contained 7×7 sub-arrays, each sub-array contained 15×15 arrays of microwells. (B) All wells within the 15×15 sub-array were numbered according to their specific position in the array. Microwells were 10 μm in diameter with a 40 μm pitch.

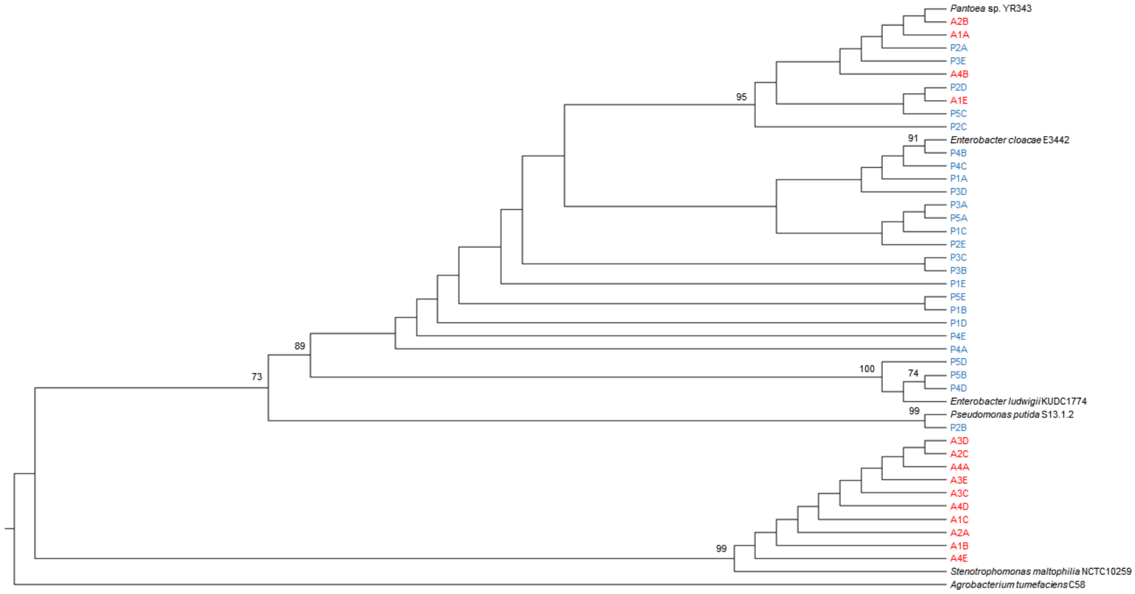

**Fig. *S2*.** Maximum likelihood phylogenetic tree based on partial sequence of the 16S rRNA (922 bp) from isolates obtained from microwells in which YR343 growth was promoted or antagonized as well as a few reference strains. The tree was constructed using the Tamura and Nei (1993) substitution model (12) with a gamma distribution (TN93 + G) in PhyML 3.0 (10). We used the Smart Model Selection (13) to select this substitution model. Bootstrap values (expressed as a percentage of 1000 replications) higher than 70% are shown at nodes. *A. tumefaciens* C58 was included as an outgroup organism.

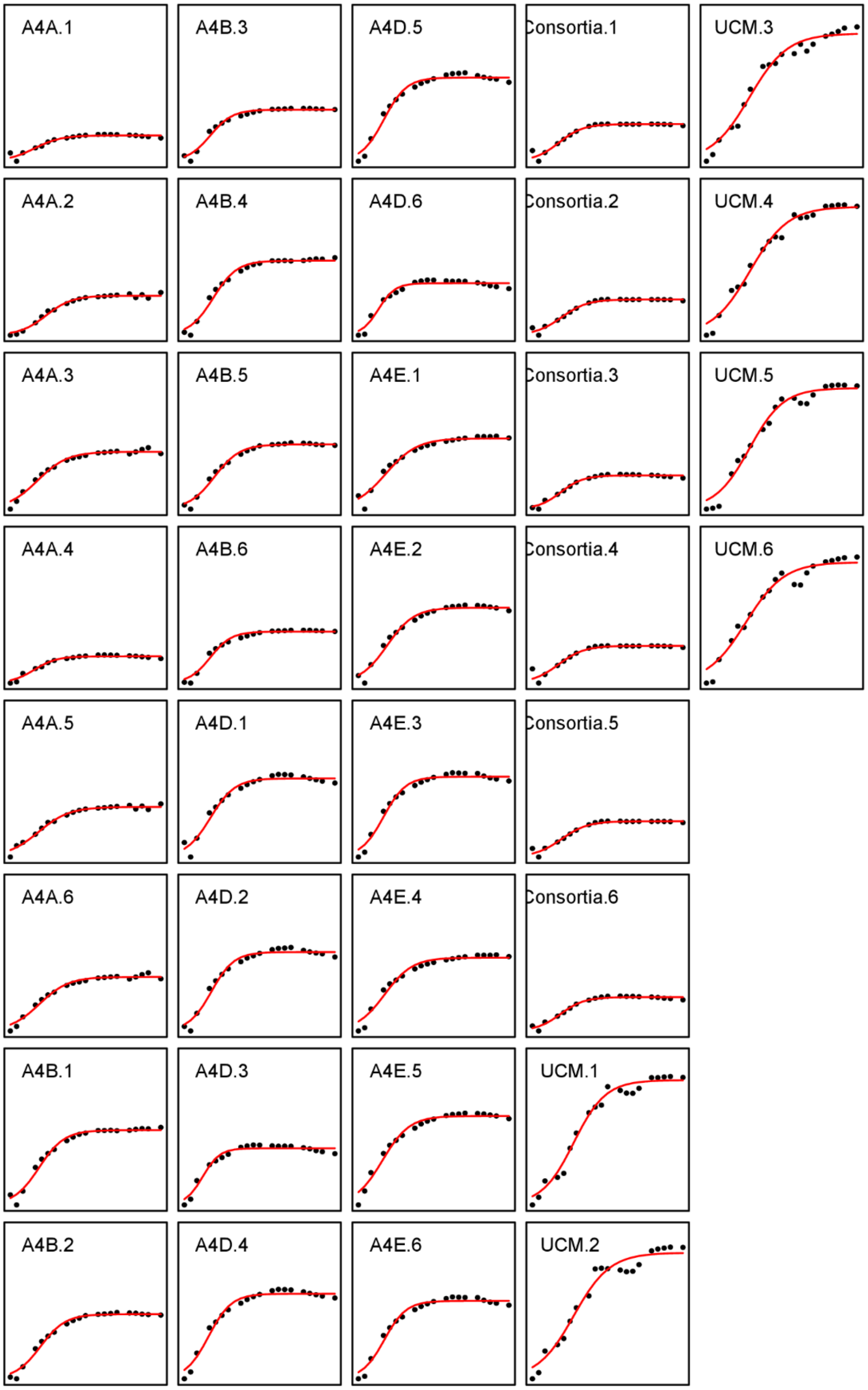

**Fig. *S3***. Growthcurver output for analysis of growth curves generated from 96 well-plate validation assays using isolates from microwells within which YR343-GFP growth was antagonized. For each condition, YR343-GFP culture was measured for a total of n=6 independent replicates. Carrying capacity, k ($OD_{600}$) and growth rate, r ($hr^{-1}$) for each isolate, isolate combination and control and were quantified (11).

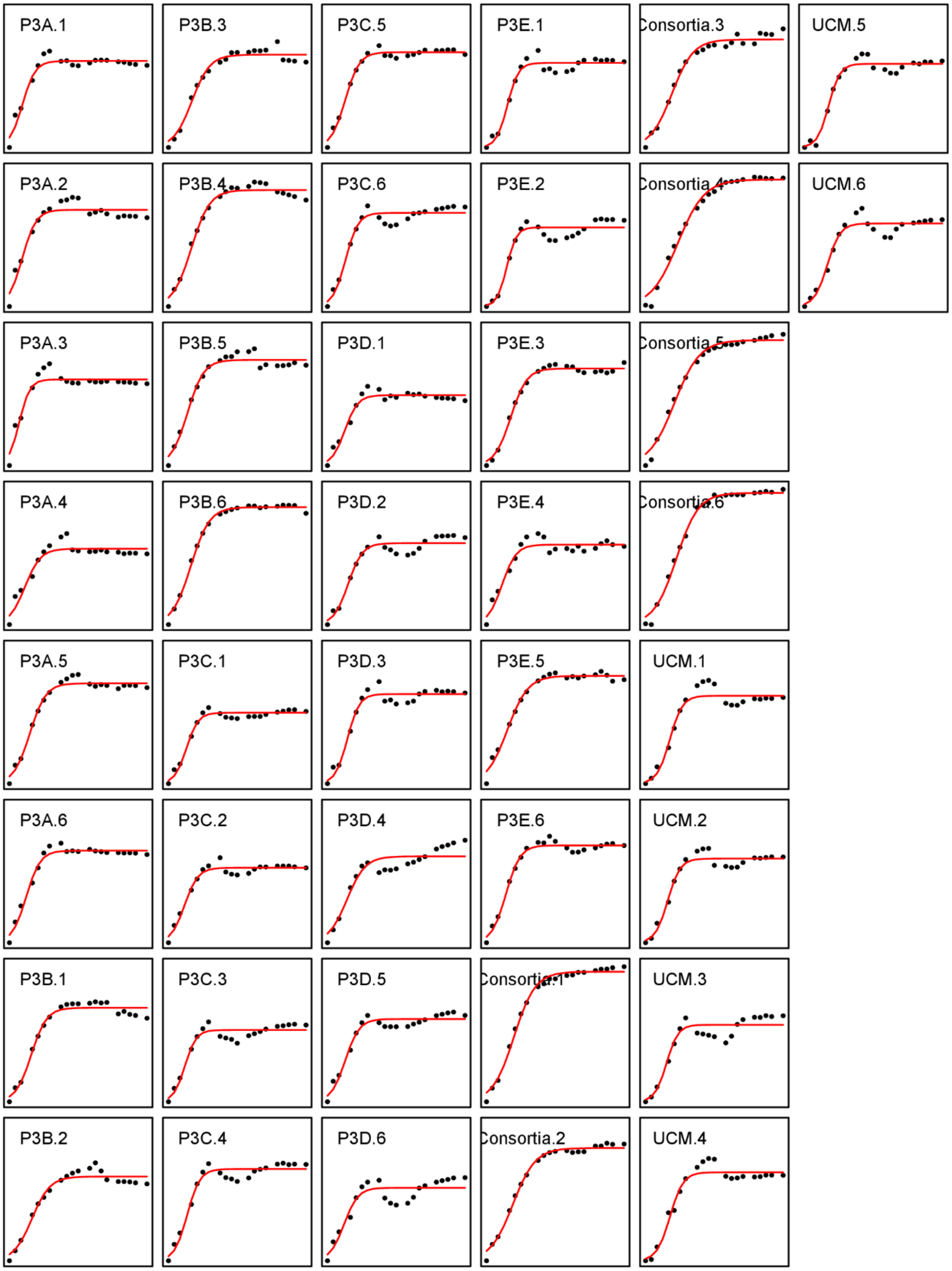

**Fig. *S4*.** Growthcurver output for analysis of growth curves generated from 96 well-plate validation assays using isolates from microwells within which YR343-GFP exhibited promoted population growth. For each condition, YR343-GFP culture was measured for a total of n=6 independent replicates. Carrying capacity, k ($OD_{600}$) and growth rate, r ($hr^{-1}$) for each isolate, isolate combination and control and were quantified (11).

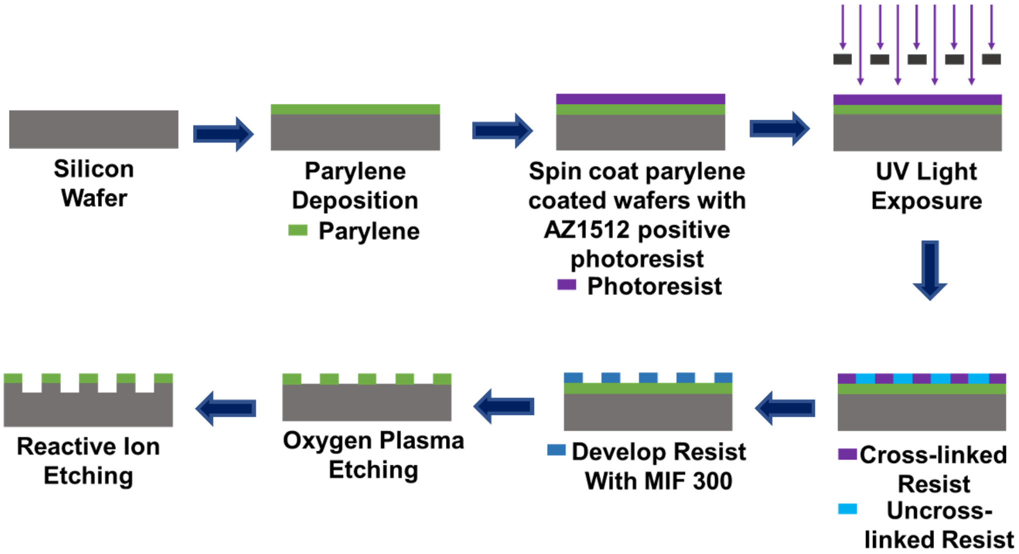

**Fig.** ***S5.*** Steps for microwell recovery array fabrication. Parylene N was deposited on top of silicon wafers. Then the parylene coated wafers were spin coated with positive photoresist AZ1512 and exposed with UV light through a photomask. The uncrosslinked photoresist was then washed by developing in MIF 300. Then Bosch etching was performed to get MRAs.

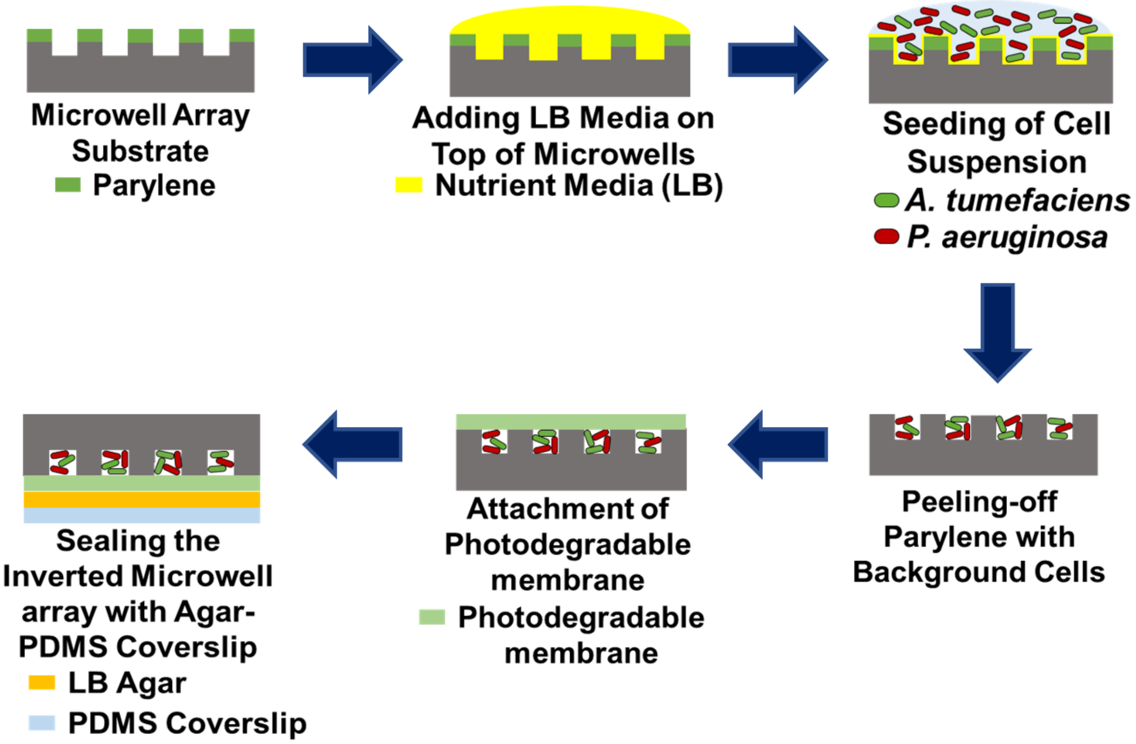

**Fig. *S6*.** Seeding and trapping of *P. aeruginosa* and *A. tumefaciens* in microwell arrays with the aid of a PDMS coverslip.

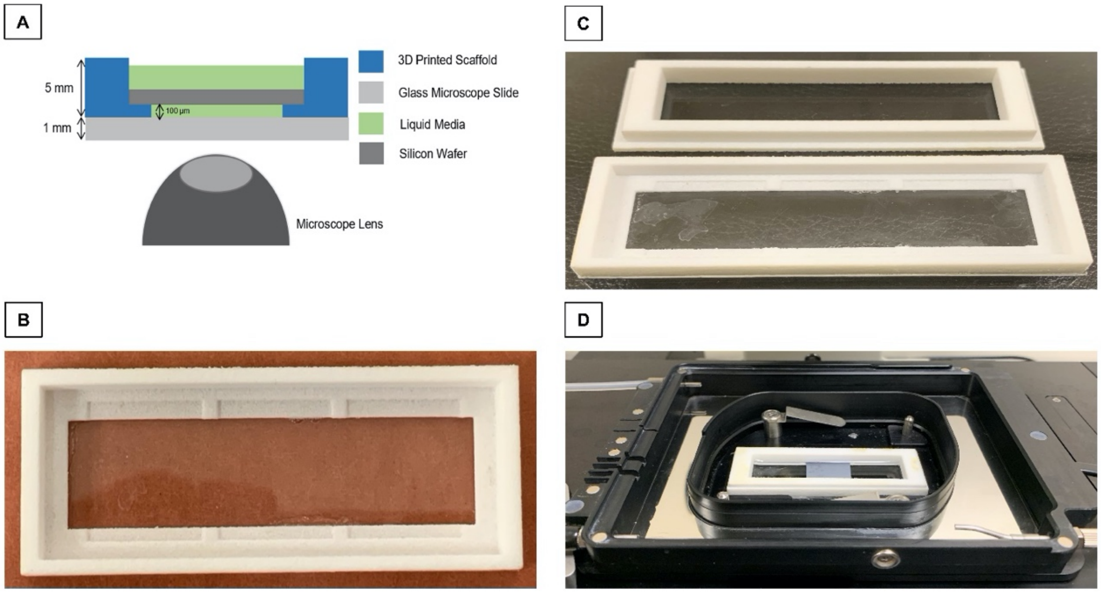

**Fig. *S7*.** (A) Cross section of the 3D printed scaffold on a glass microscope slide. (B) 3D printed Nylon scaffolds. The scaffold had 1.5×1.5cm grooves to hold up to three microwell arrays. (C) The scaffold was glued to a 75×25mm glass slide then the scaffold lid was attached on top to firmly hold the microwell in place. Liquid nutrient media was then added to fill the space between microwell array and the glass slide, keeping the microwell array fully submerged during culture. (D) The scaffold setup was placed inside a humidified live cell imaging chamber for TLFM.

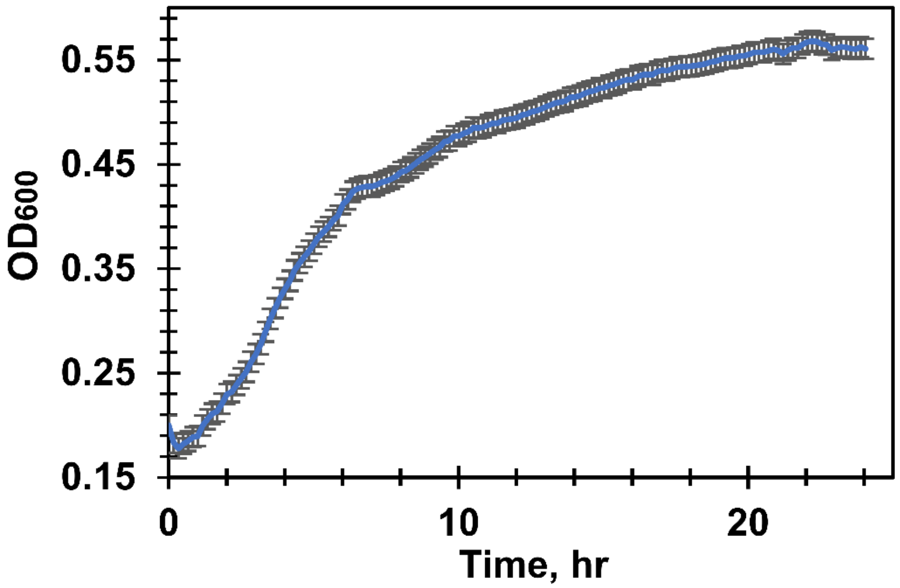

**Fig.** ***S8***. Growth curve of YR343-GFP during culture in R2A broth media at 28ºC (n=5). The lag time for YR343 growth was 2.5 hours, growth rate 0.673 $hr^{-1}$. Stationary phase was achieved after 8 hrs.

| Strain or Plasmid | Characteristics | Source or reference |
|---|---|---|
| **Strains** | | |
| *Agrobacterium tumefaciens* C58 | Wild-type strain | C. Fuqua |
| *A. tumefaciens* C58 pSRKKm-sfGFP | Wild-type strain carrying pSRKKm-sfGFP | This study |
| *Pseudomonas aeruginosa* PAO1 | Wild-type strain | ATCC |
| *P. aeruginosa* PAO1 pSRKKm-mCherry | Wild-type strain carrying pSRKKm-mCherry | This study |
| *Pantoea* sp. YR343 | Wild-type strain with a constitutively expressed chromosomal insertion of EGFP | 14 |
| *Escherichia coli S17-1 λpir* | λpir, Tra+, cloning strain | 15 |
| *E. coli S17-1 λpir* pSRKKm-sfGFP | Donor strain carrying pSRKKm-sfGFP | This study |
| *E. coli S17-1 λpir* pSRKKm-mCherry | Donor strain carrying pSRKKm-mCherry | This study |
| P1A | Isolate from microwell P1 within which YR343 exhibited promoted growth | This study |
| P1B | Isolate from microwell P1 | This study |
| P1C | Isolate from microwell P1 | This study |
| P1D | Isolate from microwell P1 | This study |
| P1E | Isolate from microwell P1 | This study |
| P2A | Isolate from microwell P2 within which YR343 exhibited promoted growth | This study |
| P2B | Isolate from microwell P2 | This study |
| P2C | Isolate from microwell P2 | This study |
| P2D | Isolate from microwell P2 | This study |
| P2E | Isolate from microwell P2 | This study |
| P3A | Isolate from microwell P3 within which YR343 exhibited promoted growth | This study |

| | | |
|---|---|---|
| P3B | Isolate from microwell P3 | This study |
| P3C | Isolate from microwell P3 | This study |
| P3D | Isolate from microwell P3 | This study |
| P3E | Isolate from microwell P3 | This study |
| P4A | Isolate from microwell P4 within which YR343 exhibited promoted growth | This study |
| P4B | Isolate from microwell P4 | This study |
| P4C | Isolate from microwell P4 | This study |
| P4D | Isolate from microwell P4 | This study |
| P4E | Isolate from microwell P4 | This study |
| P5A | Isolate from microwell P5 within which YR343 exhibited promoted growth | This study |
| P5B | Isolate from microwell P5 | This study |
| P5C | Isolate from microwell P5 | This study |
| P5D | Isolate from microwell P5 | This study |
| P5E | Isolate from microwell P5 | This study |
| A1A | Isolate from microwell A1 within which YR343 exhibited antagonized growth | This study |
| A1B | Isolate from microwell A1 | This study |
| A1C | Isolate from microwell A1 | This study |
| A1D | Isolate from microwell A1 | This study |
| A1E | Isolate from microwell A1 | This study |
| A2A | Isolate from microwell A2 within which YR343 exhibited antagonized growth | This study |
| A2B | Isolate from microwell A2 | This study |
| A2C | Isolate from microwell A2 | This study |
| A2D | Isolate from microwell A2 | This study |
| A2E | Isolate from microwell A2 | This study |
| A3A | Isolate from microwell A3 within which YR343 exhibited antagonized growth | This study |

| | | |
|---|---|---|
| A3B | Isolate from microwell A3 | This study |
| A3C | Isolate from microwell A3 | This study |
| A3D | Isolate from microwell A3 | This study |
| A3E | Isolate from microwell A3 | This study |
| A4A | Isolate from microwell A4 within which YR343 exhibited antagonized growth | This study |
| A4B | Isolate from microwell A4 | This study |
| A4C | Isolate from microwell A4 | This study |
| A4D | Isolate from microwell A4 | This study |
| A4E | Isolate from microwell A4 | This study |
| **Plasmids** | | |
| pSRKKm | Broad-host-range $P_{lac}$ expression vector; KmR | 16 |
| pSRKKm-mCherry | IPTG-inducible mCherry expression vector derived from pSRKKm; KmR | 3 |
| pSRKKm-sfGFP | IPTG-inducible GFP expression vector derived from pSRKKm; KmR | 17 |

**Table *S1*:** Bacterial strains used in this study.

| Isolate ID | p-value for Wilcoxon two-sample test | Significance of difference |
|---|---|---|
| **P3A** | 0.2876 | Not Significant |
| **P3B** | 0.0093 | Significant |
| **P3C** | 0.4051 | Not Significant |
| **P3D** | 0.0931 | Not Significant |
| **P3E** | 0.1149 | Not Significant |
| **5-member consortia** | <0.01 | Significant |

**Table *S2*:** Wilcoxon two-sample tests for differences in carrying capacities between YR343-GFP culture in conditioned versus unconditioned media from individual promoting isolates or from the 5-membered consortia.

| Isolate ID | p-value for Wilcoxon two-sample test | Significance of difference |
|---|---|---|
| **A4A** | <0.01 | Significant |
| **A4B** | <0.01 | Significant |
| **A4D** | <0.01 | Significant |
| **A4E** | <0.01 | Significant |
| **4-member consortia** | <0.01 | Significant |

**Table *S3*:** Wilcoxon two-sample tests for differences in carrying capacities between YR343-GFP culture in conditioned versus unconditioned media from individual antagonistic isolates or from the 4-membered consortia.

| Isolates ID | p-value for Wilcoxon two-sample test | Significance of difference |
|---|---|---|
| **P3A** | 0.2876 | Not Significant |
| **P3B** | <0.01 | Significant |
| **P3C** | 0.1490 | Not Significant |
| **P3D** | 0.0656 | Not Significant |
| **P3E** | <0.01 | Significant |
| **5 member consortia** | <0.01 | Significant |

**Table *S4*:** Wilcoxon two-sample tests for differences in growth rates between YR343-GFP culture in conditioned versus unconditioned media from individual promoting isolates or from the 5-membered consortia.

| Isolate ID | p-value for Wilcoxon two-sample test | Significance of difference |
|---|---|---|
| **A4A** | <0.01 | Significant |
| **A4B** | <0.01 | Significant |
| **A4D** | <0.01 | Significant |
| **A4E** | <0.01 | Significant |
| **4 member consortia** | <0.01 | Significant |

**Table *S5*:** Wilcoxon two-sample tests for differences in growth rates between YR343-GFP culture in conditioned versus unconditioned media from individual antagonistic isolates or from the 4-membered consortia.
